# In vivo colonic epithelial cell editing attenuates intestinal inflammation in mice

**DOI:** 10.1101/2025.10.08.678706

**Authors:** Hailing Zhang, Hengxing Lu, Shaolong Zhang, Xukai Hu, Qixing Wu, Zi qin Yu, Natasha Karyn Lienanto, Jin Zhang

**Author notes:** These authors contributed equally (Hailing Zhang, Hengxing Lu, Shaolong Zhang).

## Abstract

Engineered epithelial cells with enhanced efferocytosis capacity promote inflammation resolution and restore tissue homeostasis. Here, we developed a therapeutic approach to generate efferocytosis boosting epithelial cells in vivo by delivering mRNA in lipid nanoparticles (LNPs). We demonstrate that LNP-mediated delivery of mRNA enables efficient in vitro and in vivo functional editing of epithelial cells. In murine models of colitis, intraperitoneal administration of mRNA-loaded nanoparticles designed to boost efferocytosis markedly attenuated intestinal inflammation and halted disease progression. This strategy provides a proof of concept that epithelial cells can be functional engineered in situ and represents a promising therapeutic avenue for mitigating inflammatory tissue damage.

**Graphical Abstract:** 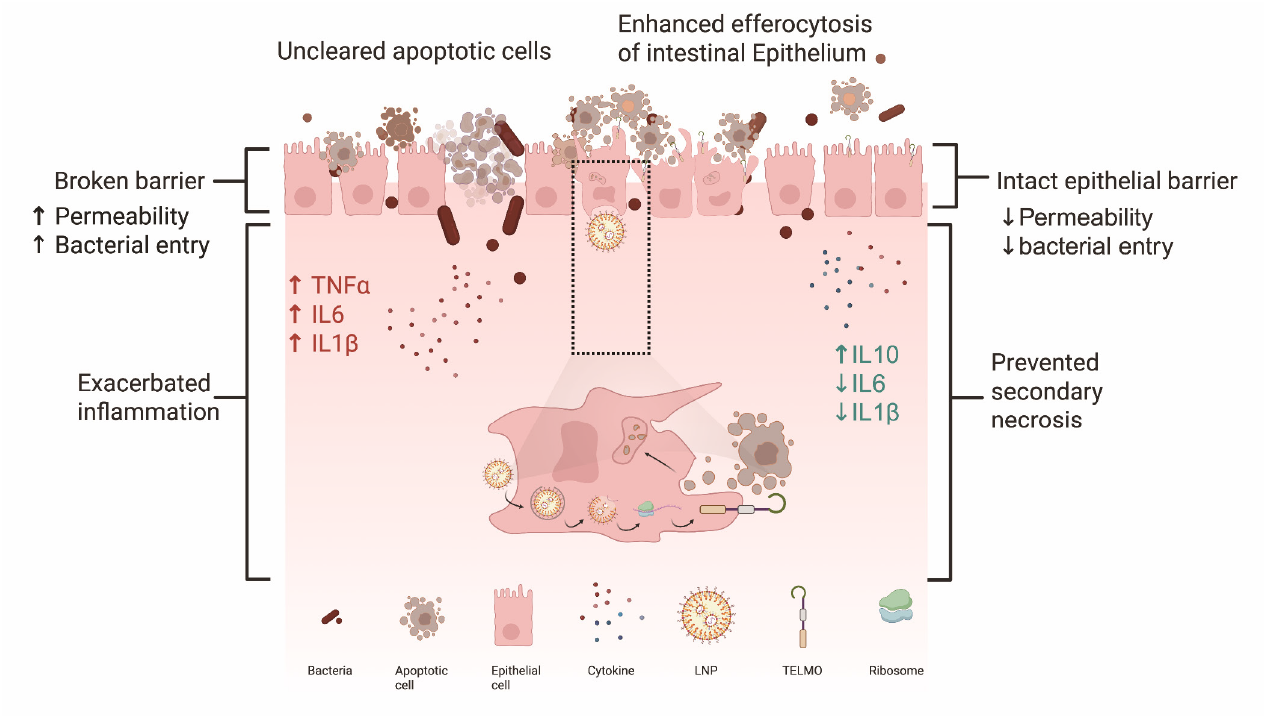

## Introduction

Phagocytes mediate the rapid clearance of apoptotic cells in a process known as efferocytosis, which is critical for tissue homeostasis and inflammation resolution^1–6^. Inadequate efferocytosis has been associated with multiple inflammatory disorders, such as atherosclerosis, systemic lupus erythematosus (SLE) and inflammatory bowel disease (IBD), contributing to pathological inflammation and tissue damage^7–10^. Sophisticated studies in transgene-overexpressing mice have demonstrated that enhancing apoptotic cell clearance can be beneficial for resolution of inflammation^11–13^. In clinical practice, editing phagocytes in situ to prompt efferocytosis may offer new opportunities for treating these diseases. In this study, we explore the use of lipid-based nanoparticles (LNPs) to deliver a chimeric efferocytic receptor (TELMO)^12^, with the goal of boosting efferocytosis as a therapeutic strategy against intestinal inflammation in mice. In a dextran sulfate sodium (DSS)-induced colitis model, intraperitoneal administration of LNPs preferentially targeted intestinal epithelial cells, resulting in robust TELMO expression and accelerated clearance of apoptotic corpses, which ultimately ameliorated inflammation and promoted tissue repair.

## Results

We generated modified mRNA encoding an efferocytosis boosting chimeric receptor, termed TELMO, which was constructed by fusing the extracellular phosphatidylserine recognition domains of TIM4 with the intracellular effector domain of ELMO1. To facilitate monitoring of TELMO expression, an enhanced green-fluorescent protein (EGFP) reporter was co-expressed via a 2A peptide (Fig. 1A). As shown in Fig. 1B and Fig. S1, the LNPs we produced demonstrated normal particle size and zeta potential values (Fig. 1B, S1). To determine whether the mRNA could be efficiently delivered into epithelial cells, we packaged the TELMO-2A-EGFP mRNA to LNPs and incubated them with HCT116 cells in vitro. Flow cytometry analysis demonstrated that LNPs were highly efficient in delivering mRNA to cultured epithelial cells, resulting in robust cell surface expression of TELMO in nearly all cells following transfection (Fig. 1C). To assess the ability of the LNP-editing cells to engulf apoptotic cells, we used staurosporine (STS)-treated Jurkat cells as apoptotic targets^14^. Successful induction of apoptosis was confirmed by Annexin V staining (Fig. S2A). The engulfment of CellTrace-labeled apoptotic cells by TELMO-expressing epithelial cells was progressively elevated over time, as shown by flow cytometry at multiple time points (Fig. 1D, Fig. S2B). This demonstrated that LNP-meditated TELMO expression significantly boosted apoptotic cell uptake. Confocal imaging also corroborated these findings, revealing more apoptotic cells localized around and within phagocytes (Fig. 1E). This observation suggests efficient internalization, prompting further investigation into the ensuing intracellular processing dynamics. To investigate the intracellular processing of apoptotic cells after TELMO-mediated efferocytosis, we performed a corpse acidification assay using a dye-quenching approach (Fig. 1F). TELMO expression significantly enhanced acidification, TELMO expression significantly enhanced the acidification process, as shown by a reduction in mean fluorescence intensity (MFI), indicating accelerated apoptotic cargo processing (Fig. 1G, S3A). To further validate this observation, we utilized a pH-sensitive reporter dye that fluoresces specifically in acidic compartments. TELMO not only increased the percentage of phagocytes that trafficked apoptotic corpses to acidic organelles (Fig. 1H, S3B), but also elevated the acidified cargo intensity per cell (Fig. 1I, S3C). Additionally, confocal microscopy confirmed that TELMO promotes both efficient internalization and processing of apoptotic cells (Fig. 1J).

**Figure 1.**
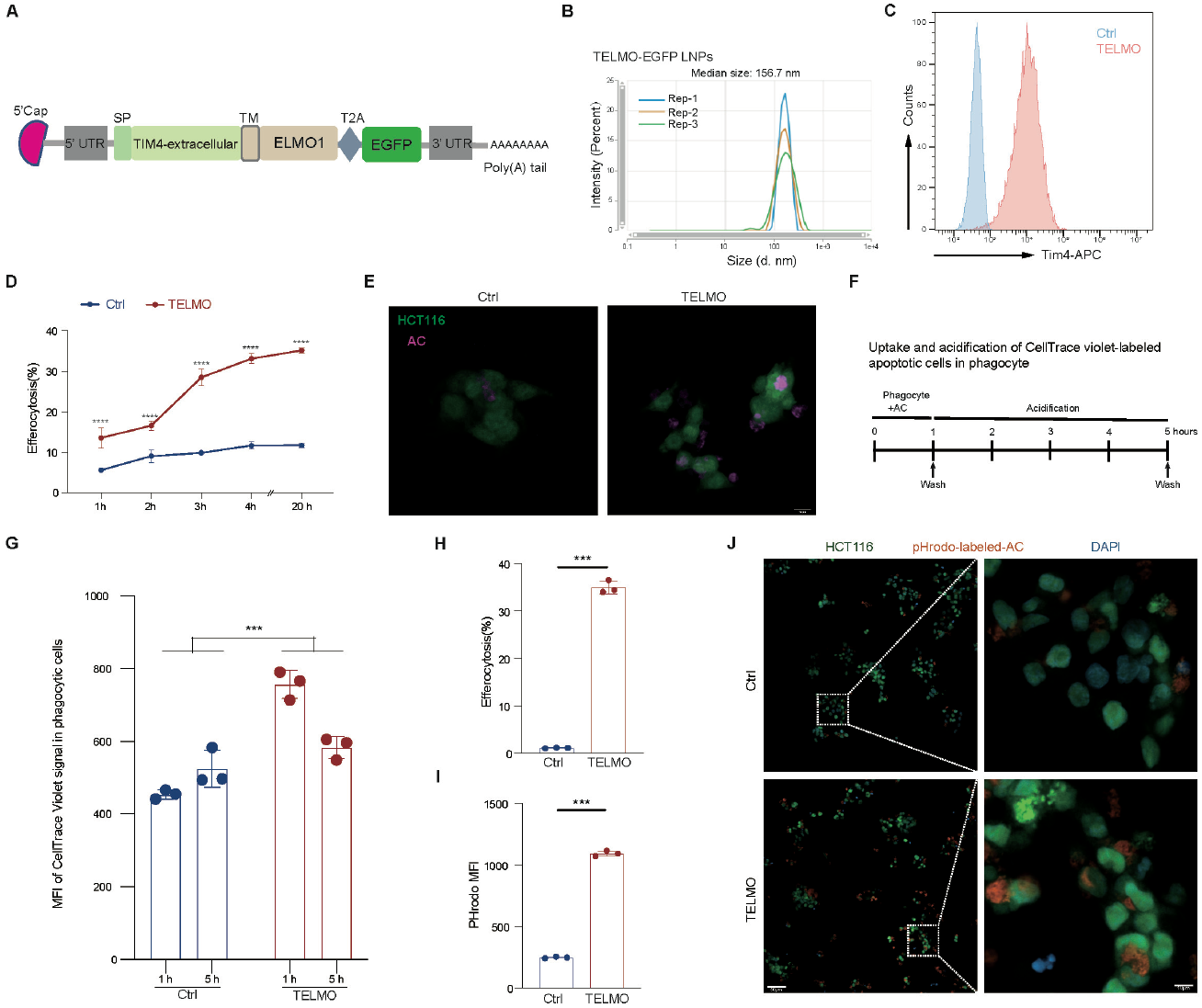
LNP-mediated TELMO expression boosts phagocytosis of apoptotic cells. (A) Schematic diagram of a TELMO expression construct used for lipid nanoparticle (LNP)-mediated mRNA delivery. SP, signal peptide; TM, transmembrane domain. (B) Particle diameter distribution of TELMO-EGFP LNPs. (C) Flow cytometry analysis of TELMO surface expression in HCT116 cells 24 h after TELMO mRNA-LNP transfection. (D) Quantification of efferocytosis kinetics over time (1-20 hours). (E) Confocal microscopy images depicting TELMO-expressing HCT116 cells engulfing apoptotic Jurkat cells. Scale bar: 10 µm. (F) Schematic for the apoptotic corpse acidification assay. (G) Quantification of acidification levels (MFI reduction) in phagocytosed cargo (n=3). (H) Quantification of efferocytosis-positive phagocytes. (I) Quantification of per-cell acidified cargo intensity. (J) Confocal images confirming apoptotic cell internalization and processing of apoptotic cells.

We next examined the ability of LNP mRNA to modify epithelial cells in vivo. TELMO-EGFP-encoding LNPs were administered by intraperitoneal injection in mice, and colon tissues were collected after 24 hours for analysis. EGFP expression was observed in colonic epithelial cells by fluorescence microscopy, indicating effective LNP-mediated delivery and functional expression of the engineered constructs (Fig. 2A). Further flow cytometry analysis demonstrated that TELMO was expressed in approximately 50% of EpCAM-positive cells, thus confirming the highly efficient engineering of intestinal epithelium via LNP-mediated mRNA delivery (Fig. 2B, S4). Encouraged by the efficient in vivo engineering of colonic epithelial cells, we next evaluated the therapeutic potential of TELMO mRNA delivery in a mouse model of colitis. TELMO-LNPs or, as a control, GFP-LNPs were intraperitoneally (i.p.) injected into DSS-exposed C57BL/6 mice according to the procedure outlined depicted in Figure 2C. The colitic mice received LNP treatments on days 0 and 3. Clinical signs of colitis, including body weight change, stool consistency, and rectal bleeding, were monitored daily. On day 10, colon length and histology were evaluated. We found that TELMO-LNP treatment significantly mitigated the clinical manifestations of DSS-induced colitis. Mice receiving TELMO-LNPs showed significantly reduced disease severity, as evidenced by attenuated body weight loss (Fig. S5A), improved stool consistency (Fig. S5B), and less rectal bleeding (Fig. S5C), resulting in lower Disease Activity Index (DAI) scores (Fig. 2D and S5D). Moreover, intestinal epithelial permeability was analyzed by FITC-dextran leakage assay. The serum level of FITC-Dextran was significantly increased in control mice, but this increase was markedly attenuated by TELMO-LNP treatment (Fig. 2E). Consistent with the symptom improvement, TELMO-LNPs also alleviated colon shortening (Fig. 2F, 2G), indicating reduced tissue damage and inflammation at the gross anatomical level. This result suggests that TELMO-LNPs helped restore intestinal barrier integrity, thereby preventing bacterial translocation and subsequent systemic inflammation. Corresponding to these functional improvements, histological examination also confirmed preservation of tissue architecture at the microscopic level. Compared to vehicle controls, TELMO-LNP-treated mice exhibited more intact crypt structures, diminished mucosal erosion, and reduced inflammatory cell infiltration (Fig. 2H, 2I). TUNEL staining further demonstrated minimal epithelial apoptotic cell death following TELMO-LNP treatment (Fig. 3A, 3B), suggesting that tissue integrity was preserved through active cytoprotection rather than merely delayed inflammatory damage. To evaluate the level of tissue inflammation, we performed qPCR analysis on colonic tissues. The results showed that the expression of pro-inflammatory cytokines (TNF-α, IL-1β, and IL-6) and chemokine-related genes (CXCL9 and CCL-2) was significantly downregulated in the treatment group. In contrast, the anti-inflammatory gene IL-10 was markedly upregulated (Fig. 3C-3H). Collectively, these results suggest that in vivo editing of colonic epithelial cells via TELMO-LNPs not only ameliorates clinical symptoms and preserves tissue architecture, but also reshapes the inflammatory microenvironment toward a more anti-inflammatory state.

**Figure 2.**
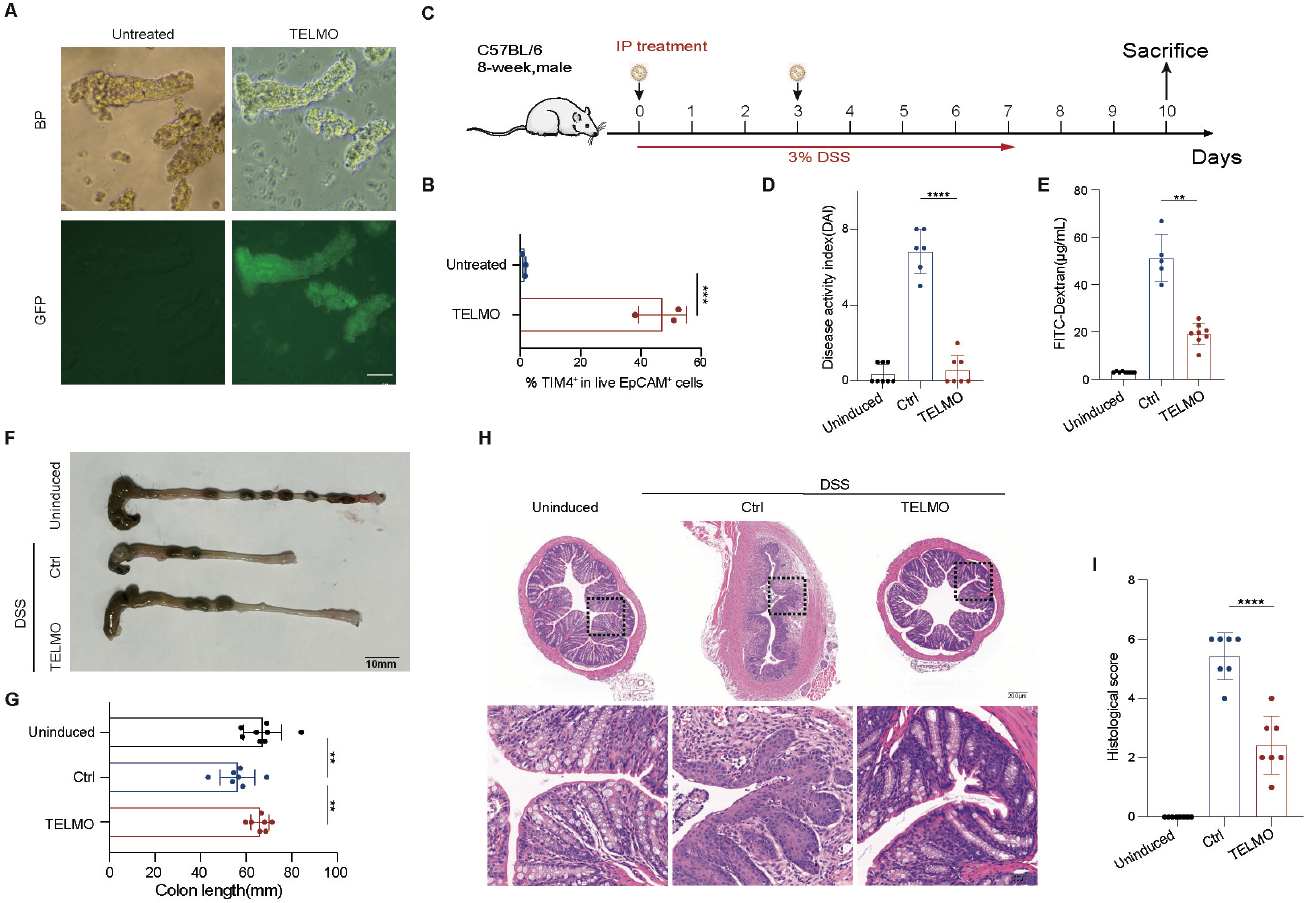
TELMO-LNPs attenuates DSS-induced colitis. (A) Representative fluorescence microscopy images showing GFP expression in colonic epithelial cells 24 hours after intraperitoneal administration of TELMO-GFP LNPs. Both brightfield (BF) and GFP channels are shown for untreated and TELMO-LNP–treated groups. Scale bar: 50 µm. (B) Quantification of TELMO expression efficiency in EpCAM+ colonic epithelial cells(n=3). (C) Summary of the experimental protocol for DSS-induced colitis model (n=7-10). (D) Disease activity index (DAI) scores for colitis mice. (E) Serum FITC–dextran concentration as an index of intestinal permeability. (F, G) Representative images and quantification of colon length at sacrifice. (H) Representative Hematoxylin and eosin (H&E) staining of distal colon sections. Scale bars: 200 µm (overview), 50 µm (zoom). (I) Histologic score derived from the H&E-stained sections.

**Figure 3.**
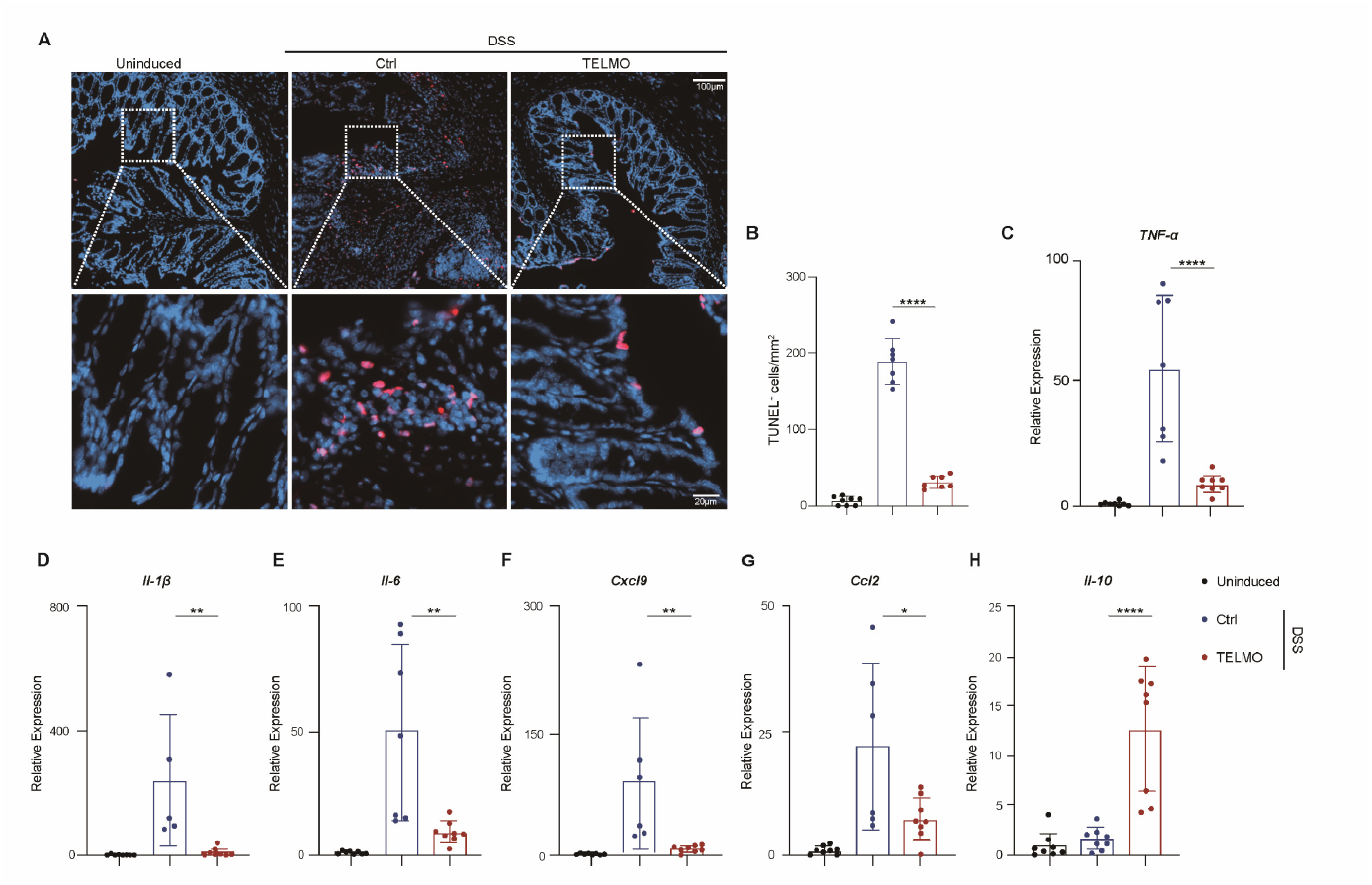
TELMO-LNPs reduce the apoptotic cell burden in tissues and alleviate tissue inflammation. (A) TUNEL staining of colon sections. 100 µm (overview), 20 µm (zoom). (B) Quantitative analysis of TUNEL staining. (C-H) Colonic mRNA expression of pro-inflammatory cytokines (TNF-α, IL-1β, and IL-6), chemokine-related genes (CXCL9 and CCL-2) and the anti-inflammatory gene IL-10. Data are presented as mean (SEM); significance was determined by unpaired Student’s two-tailed t-test, one-way ANOVA, or two-way ANOVA according to test requirements, *P<0.05, **P<0.01, ***P<0.001.

## Discussion

Gut inflammatory disorders, such as ulcerative colitis (UC) and IBD affect millions of people worldwide. Conventional therapeutic approaches are primarily ameliorative and often fail to address the underlying etiology of these diseases^15^. Colonic epithelial cells constitute the physical barrier and are fundamentally required for maintaining its integrity. Their role in clearing apoptotic neighbors is critical for the resolution of intestinal inflammation^11,16^. The rapid, potent, and transient expression of mRNA-encoded proteins, which avoids the risk of detrimental genomic integration, makes them an attractive therapeutic platform^17^. In recent years, in vitro transcribed mRNA has shown promise for treating a broad spectrum of diseases, including cancer, fibrosis, and autoimmune diseases^18–22^. Our present study demonstrated that LNP-mediated mRNA delivery effectively edits colonic epithelial cells, enhances efferocytosis in a local tissue environment, and confers therapeutic benefits, providing a convenient and effective strategy for treating inflammatory diseases via efferocytic reprogramming of phagocytes in situ. In our observations, direct intraperitoneal administration of LNPs achieved efficient editing in intestinal epithelial cells, likely attributable to their preferential uptake by phagocytic cells. In contrast, other immune cells in the intestinal tissue, such as T cells, B cells, and macrophages, either exhibited low abundance or minimal editing efficiency, suggesting that epithelial cells serve as the principal mediators of the therapeutic effect. Further studies are needed to optimize the dosing regimen, LNP formulation, and targeting strategies to improve therapeutic outcomes and limit potential toxicities.

## Acknowledgements

We thank the Core Facilities of Liangzhu Laboratory. This work was funded by the National Natural Science Youth Foundation of China (No. 82202039).

## Author contributions

J.Z, H.L, and H.Z conceived and designed the experiments. H.Z, H.L, and S.Z performed mainly experiments. H.Z, S.Z and Q.W conducted the mRNA and LNPs production. X.K, Z.Y and N.L participated in animal experiments. H.Z, H.L, and J.Z wrote the paper. All authors contributed to the paper and approved its final version.

## Supplementary information

### Materials and methods

#### Cell culture

The HCT116 cells were cultured in McCoy’s 5A medium supplemented with 10% fetal bovine serum (FBS) and 1% penicillin-streptomycin (P/S) solution. Jurkat T cells were maintained in culture using RPMI 1640 supplemented with 10% FBS and 1% P/S. The cells were incubated at 37 °C in a 5% CO2 humidified atmosphere.

#### RNA Synthesis and Lipid Nanoparticle Encapsulation

The TELMO construct, as previously described^1^, consists of the extracellular phosphatidylserine recognition domains of the efferocytic receptor TIM4 and a specific signaling domain of ELMO (Fig. S1). The sequence was cloned into a plasmid encompassing a T7 promoter, 5’ and 3’ UTR elements, and a poly(A) tail. Modified mRNA was synthesized using in vitro transcription with T7 RNA polymerase. The transcription reaction was assembled in a 20 µL volume containing 1× transcription buffer, 10 mM each of ATP, CTP, GTP, and N1-methylpseudouridine-5’-triphosphate (N1-Me-pUTP), 10 mM 5’-capping analog (CAP GAG 3’OMe), 1 µg template DNA, and T7 RNA polymerase mix. The reaction was incubated at 37°C for 4-8 hours. Template DNA was removed by treatment with RNase-free DNase I at 37°C for 15 minutes. RNA was purified using lithium chloride precipitation. Briefly, the transcription reaction was diluted with 30 µL RNase-free water and 30 µL of 7.5 M LiCl, then precipitated overnight at -20°C. The precipitate was collected by centrifugation at maximum speed for 15 minutes at 4°C, washed with ice-cold 70% ethanol, air-dried, and resuspended in 50 µL RNase-free water to achieve a final concentration of approximately 1 µg/µL. Purified mRNA was stored at -80°C until use. The purified RNA was encapsulated into LNPs by rapidly mixing it with an ethanol lipid mixture containing separable cationic lipids, phospholipids, cholesterol, and PEG-lipid at an acidic pH.

#### Apoptosis induction and assessment

For induction of apoptosis, human Jurkat T cells, resuspended in RPMI with 1% BSA, were treated with 1 µM staurosporine (STS; MCE) and incubated for 4 h at 37 °C and 5% CO_2_. Then apoptotic cells were stained with annexin V-FITC (BioLegend) for 15 min at room temperature in annexin V binding buffer and Propidium Iodide (PI; Yeasen) was added 5 minutes prior to running.

#### Efferocytosis assay

The target apoptotic cells were stained with CellTrace Violet or pHrodo Red (Thermo Fisher Scientific) before use in the engulfment assays. HCT116 cells were plated in a 24-well plate, and co-incubated with targets at a 1:10 phagocyte to target ratio for the indicated times. Unbound/unengulfed targets were then washed away with PBS. After the indicated incubation times, cells were harvested for flow cytometric analysis or fixed with DAPI nuclear stain (MCE) for fluorescence microscopy examination.

#### DSS-induced colitis model

Mice (C57BL6, 8∼10 weeks old) received 3% dextran sulfate sodium (DSS; MP Biomedicals) in drinking water for the indicated days, subsequently replaced with regular drinking water until experimental termination. Changes in body weight and disease severity index were monitored daily. Disease severity was quantified using the Disease Activity Index (DAI) scoring system incorporating weight loss, occult blood, and stool consistency, as previously described^2,3^. The total scores were calculated as follows: weight loss((0: < 1 %, 1: 1∼5 %, 2: 6∼10 %, 3: 11∼20 %, 4: > 20 %), stool blood ((0: negative, 1:positive Hemoccult;2: visible blood traces in stool, 3: gross rectal bleeding), and stool consistency (0: normal;1: semi-formed stools that did not adhere to the anus;2: pasty semi-formed stool that adhered to the anus; 3: liquid stools that adhered to the anus).

#### Fluorescein isothiocyanate (FITC) labeled dextran permeability assay

Intestinal permeability was assessed through the oral FITC-dextran tracer administration. Mice underwent 4-hour food restriction before intragastric delivery of FITC-labeled dextran (4000 Da, 0.2 mg/g bodyweight, #FD150S; Sigma-Aldrich) suspended in 100 µL phosphate-buffered saline. After 4 h, mouse blood was collected to obtain the hemolysis-free serum for fluorescence intensity detection using the Tecan spark multimode microplate reader (excitation: 488 nm, emission: 520 nm). FITC-dextran serum concentrations were determined against standard dilution curves prepared in PBS buffer.

#### Histological Analysis

Distal colon segments (0.5 cm from anus) were fixed, paraffin-embedded, and sectioned (4 µm) for hematoxylin and eosin (H&E) staining. Histological scores of colitis were assessed based on an established scoring criteria.^3^ Briefly, inflammation severity was scored as: 0, normal (within normal limits); 1, mild (focal inflammation limited to lamina propria); 2, moderate (multifocal or locally extensive inflammation extending to submucosa); 3, severe (transmural inflammation with ulcers affecting >20 crypts). Ulceration was scored as: 0, normal (no ulcers); 1, mild (1-2 ulcers affecting ≤20 crypts); 2, moderate (3-4 ulcers affecting 20-40 crypts); 3, severe (>4 ulcers or >40 crypts affected). Apoptotic cell detection was performed using TUNEL methodology following manufacturer’s protocol (YSFluorTM 640; Yeasen).

#### Quantitative RT-PCR

Total RNA from the colons extracted using the FastPure Complex Tissue/Cell Total RNA Isolation Kit (Vazyme). Then, cDNA was generated using the EasyScript One-Step gDNA Removal and cDNA Synthesis SuperMix kit (TransGen Biotech). Quantitative PCR was then performed with a ChamQ Blue Universal SYBR qPCR Master Mix Kit (Vazyme) to profile the gene expressions, using histone gene *Gapdh* as the standard control. All qPCR primers (5′→3′) were designed using Primer-BLAST (NCBI) and validated experimentally. Primer sequences are provided in Supplementary Table1.

## supplementary figures

**Supplementary Figure. S1.**
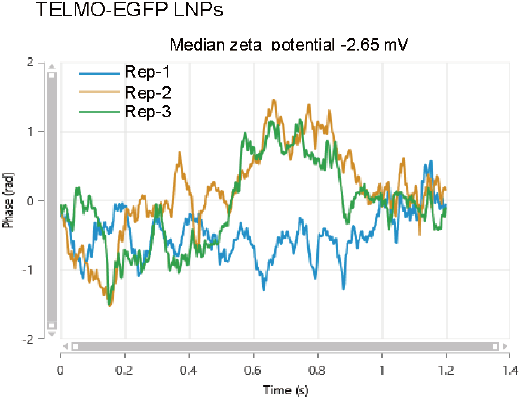
zeta potentials of TELMO-EGFP LNPs.

**Supplementary Figure. S2.**
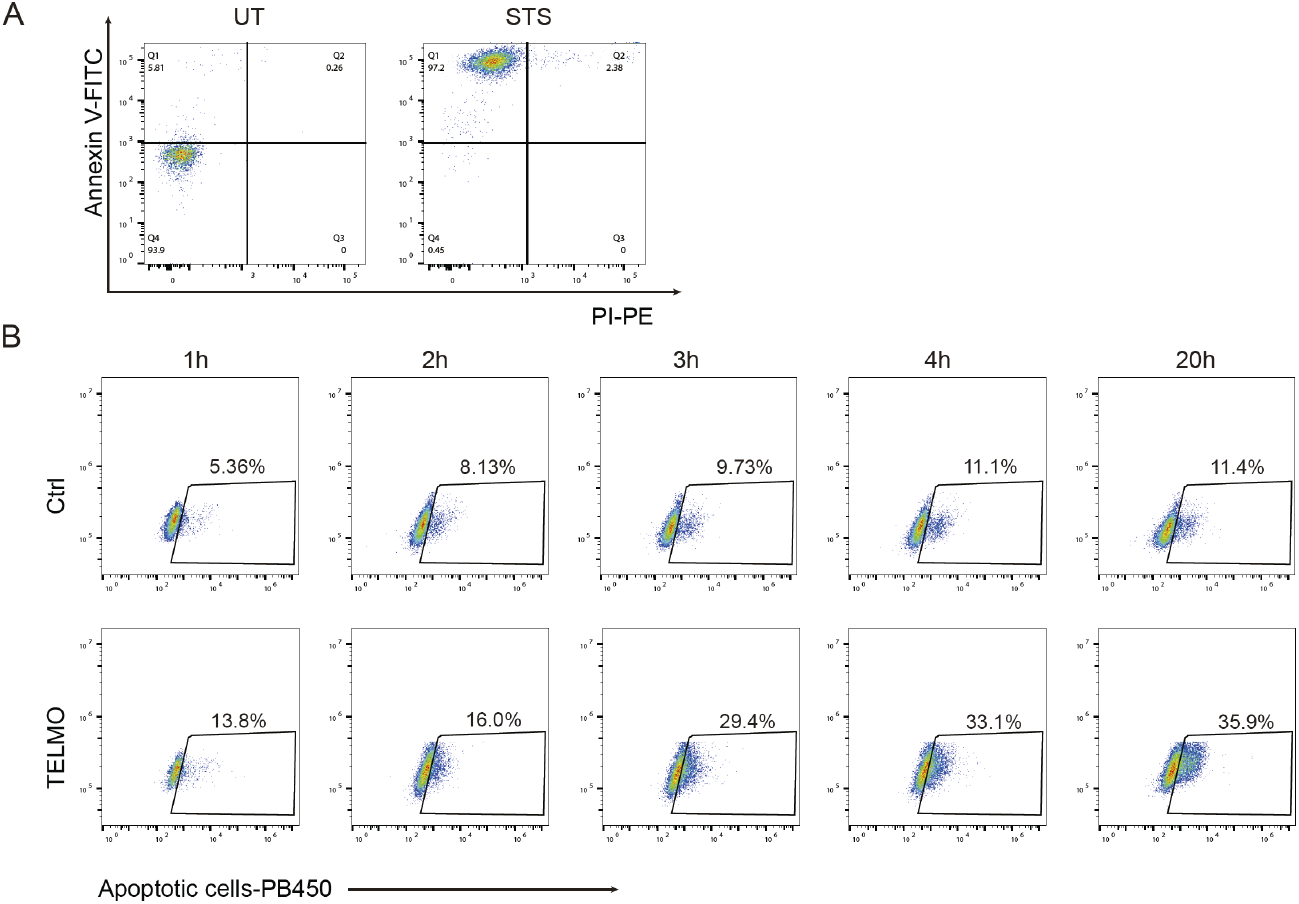
Apoptosis validation and time-course analysis of apoptotic cell uptake by TELMO-expressinq cells. (A) Flow cytometric analysis of early apoptotic Jurkat cells following 4-hour staurosporine (STS) treatmenVdetected by Annexin V and propidiumjodide (PI) double staining. (B) Representative flow cytometry plots showing uptake of CellTrace Violet-labeled apoptotic Jurkat cells.by TELMO-expressinq HCT116 cells over 1-20 h co-culture(n=3).

**Supplementary Figure. S3.**
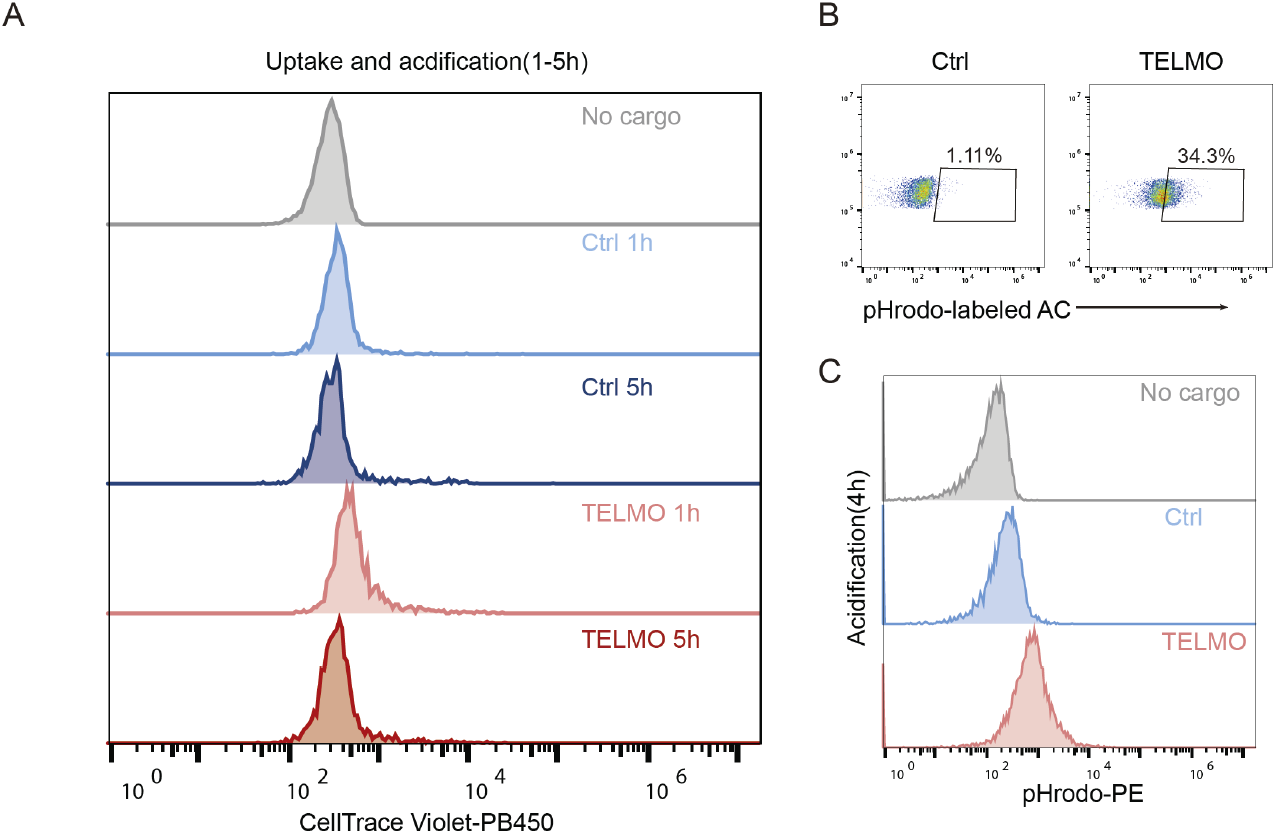
Phagolysosomal acidification in TELMO phagocytes. (A) Flow cytometry analysis of acidification levels in phagocytosed cargo, measured as mean fluorescence intensity (MFI) reduction. (B, C) Flow cytometry evaluation of acidic compartment trafficking using a pH-sensitive reporter dye: proportion of pHrodo^+^ phagocytes (B) and per-cell cargo intensity (C) (n=3).

**Supplementary Figure. S4.**
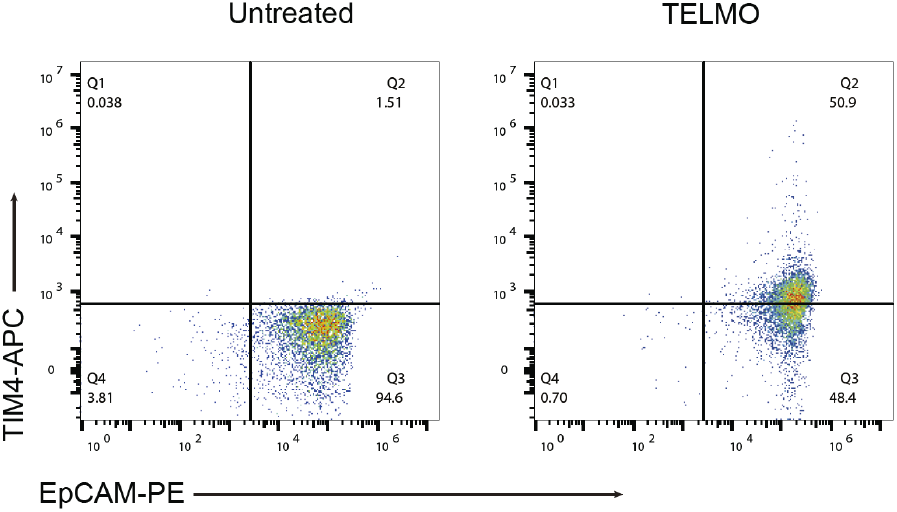
TELMO-LNP delivery to colonic epithelial cells. Flow cytometry plots showing TELMO expression in EpCAM+ colonic epithelial cells following intraperitoneal TELMO-LNP administration(n=3).

**Supplementary Figure. S5.**
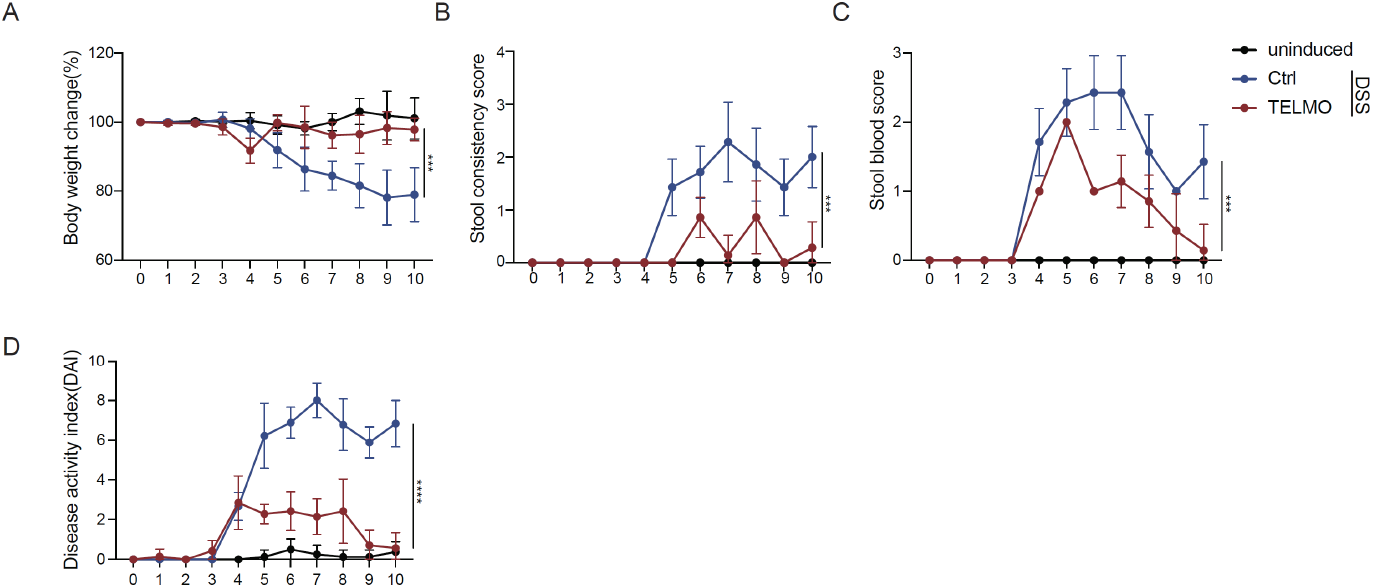
Therapeutic efficacy of TELMO-LNP treatment in DSS-induced colitis. Clinical assessment of dextran sulfate sodium (DSS)-induced colitis in vehicle-treated control and TELMO-LNP–treated mice (n = 7–8 mice per group). Parameters include: body weight change from baseline(A), stool consistency score (B), rectal bleeding score (C), and (D) composite Disease Activity Index (DAI) calculated from the sum of individual clinical scores. Data represent mean ± SEM; significance was determined by two-way ANOVA, ∗∗∗p < 0.001, ∗∗∗p < 0.0001.

**Supplementary Table 1.**
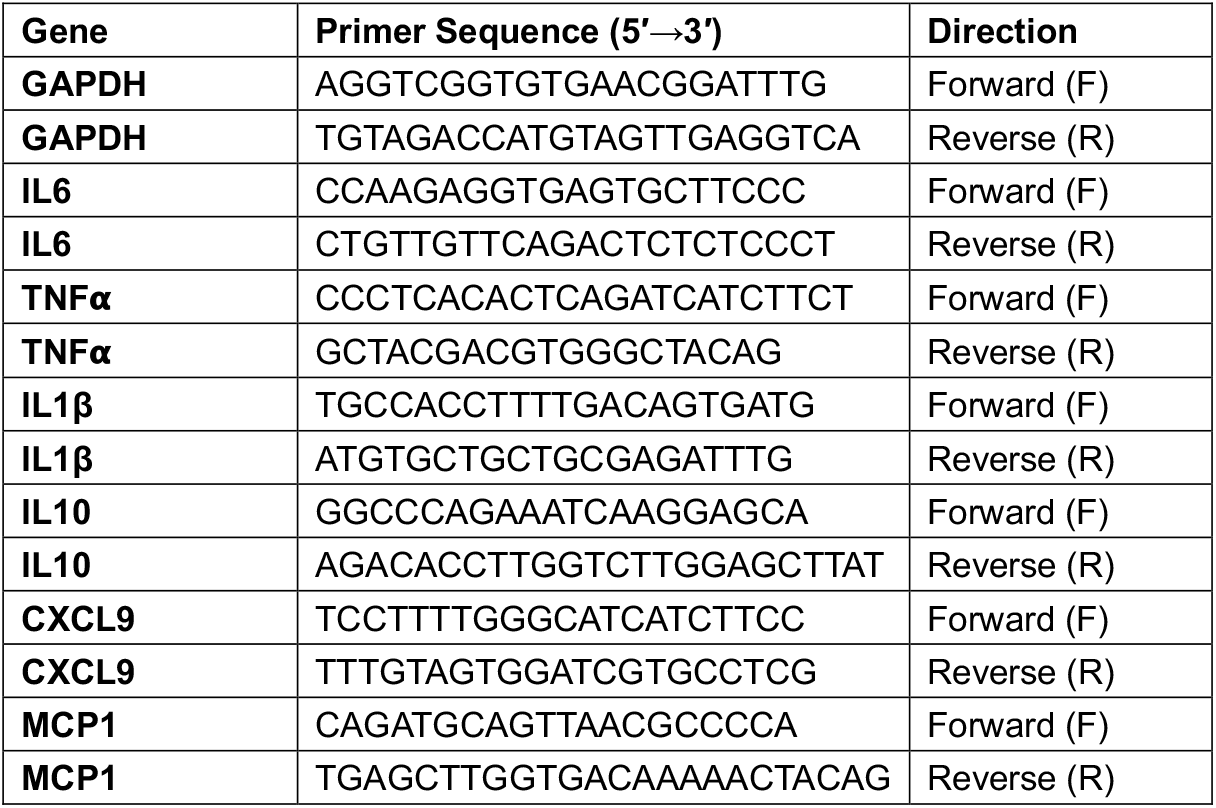
List of primers used for quantitative real-time PCR (qPCR).

